# IL-11 is a therapeutic target in idiopathic pulmonary fibrosis

**DOI:** 10.1101/336537

**Authors:** Benjamin Ng, Jinrui Dong, Sivakumar Viswanathan, Giuseppe D’Agostino, Anissa A. Widjaja, Wei-Wen Lim, Nicole SJ. Ko, Jessie Tan, Sonia P. Chothani, Benjamin Huang, Chen Xie, Ann-Marie Chacko, Nuno Guimarães-Camboa, Sylvia M. Evans, Adam J. Byrne, Toby M. Maher, Jiurong Liang, Paul W. Noble, Sebastian Schafer, Stuart A. Cook

**Affiliations:** National Heart Centre Singapore, Singapore.; Duke-National University of Singapore Medical School, Singapore.; Skaggs School of Pharmacy and Pharmaceutical Sciences, University of California San Diego, La Jolla, USA.; Department of Medicine, University of California at San Diego, La Jolla, USA.; Department of Pharmacology, University of California at San Diego, La Jolla, USA.; Fibrosis Research Group, Inflammation, Repair and Development Section, National Heart and Lung Institute, Imperial College, London, UK.; NIHR Biomedical Research Unit, Royal Brompton Hospital, London, UK.; Department of Medicine, Cedars-Sinai Medical Center, Los Angeles, California, USA.; Women’s Guild Lung Institute, Cedars-Sinai Medical Center, Los Angeles, California, USA.; National Heart and Lung Institute, Imperial College, London, UK.; MRC-London Institute of Medical Sciences, Hammersmith Hospital Campus, London, UK.

## Abstract

Idiopathic pulmonary fibrosis (IPF) remains a progressive disease despite best medical management. We previously identified IL-11 as a critical factor for cardiovascular fibrosis and examine here its role in pulmonary fibrosis. *IL-11* is consistently upregulated in IPF genomic datasets, which we confirmed by histology. Pulmonary fibroblasts stimulated with IL-11 transform into invasive myofibroblasts whereas fibroblasts from *Il11ra* deleted mice did not respond to pro-fibrotic stimuli. In the mouse, injection of recombinant Il-11 or fibroblast-specific expression of Il-11 caused pulmonary fibrosis. We then generated a neutralising IL-11 binding antibody that blocks lung fibroblast activation across species. In a mouse model of IPF, anti-IL-11 therapy attenuated lung fibrosis and specifically blocked Erk activation. These data prioritise IL-11 as an accessible drug target in IPF.

**One Sentence Summary:** Non-canonical IL-11 signalling is a central hallmark of idiopathic pulmonary fibrosis and represents a novel target for antibody therapies.

## Main Text

Idiopathic pulmonary fibrosis (IPF) is a progressive fibrotic lung disease that is characterised by epithelial injury and activation of invasive fibroblasts that deposit and remodel extracellular matrix (ECM) destroying tissue integrity (*1*, *2*). Aging, genetic and environmental factors are known to trigger IPF but the underlying pathological mechanisms remain poorly characterised (*2*). Anti-inflammatory agents do not improve clinical outcomes (*3*–*5*), emphasizing the central role of fibroblast biology in IPF. Two anti-fibrotic drugs, nintedanib (*6*) and pirfenidone (*7*), are approved for the treatment of IPF but have notable toxicities (*2*) and despite best medical practice IPF remains a fatal disease and an unmet clinical need. Unfortunately recent clinical trials in IPF have not shown efficacy and new approaches based on a better understanding of disease biology are needed (*8*).

Transforming growth factor-beta 1 (TGFβ1) is considered the principal pro-fibrotic cytokine and plays an important role in lung fibrosis (*9*), but TGFβ1-targeting therapies have marked side effects due to the pleiotropic roles of TGFβ1 in diverse cell types (*8*, *10*). We recently screened for fibroblast-specific mediators of TGFβ1 activity in human cardiac fibroblasts and identified interleukin-11 (IL-11) as a crucial fibroblast-specific factor for cardiovascular fibrosis (*11*). In the lung there is conflicting literature for IL-11 that suggests it may either promote (*12*) or protect (*13*, *14*) against lung damage, inflammation and/or fibrosis. Likely because of these discrepancies, the study of *IL-11* biology in the lung has not progressed over the past decades. In recent genome wide studies *IL-11* was found to be the most upregulated gene in fibroblasts from patients with IPF (*15*) but this chance observation was not explored further. Here we examined the hypothesis that IL-11 may be important for lung fibroblast activation and the pathobiology of IPF.

## Results

### IL-11 is upregulated in the IPF lung

To better understand *IL-11* expression in human pulmonary fibrosis, we integrated three separate genome wide RNA expression datasets in healthy and IPF lung tissue (*16*–*18*). Raw sequencing data was reprocessed and combined using a standardised pipeline (see Methods). *IL-11* expression was consistently increased in IPF in all studies (Fig. 1, A to C). When studies were combined, and after adjusting for confounding effects, we confirmed further that *IL-11* transcripts are highly upregulated (P_adj._= 3.88e-11) in the lung of IPF patients (Fig. 1D and fig. S1, A and B). We then performed immunostaining for IL-11 and alpha smooth muscle actin (ACTA2), a marker for myofibroblasts, in lung sections from healthy individuals and IPF patients. In keeping with the RNA-based analyses, IL-11 protein was lowly expressed in normal lung tissue but markedly elevated in IPF lung samples along with ACTA2 (Fig. 1E). IL-11 in the IPF lung is most likely secreted by fibroblasts, since *IL-11* RNA is very markedly upregulated in lung fibroblasts isolated from patients with systemic sclerosis or IPF (data reanalysed from (*15*)) (Fig. 1F).

**Fig. 1.**
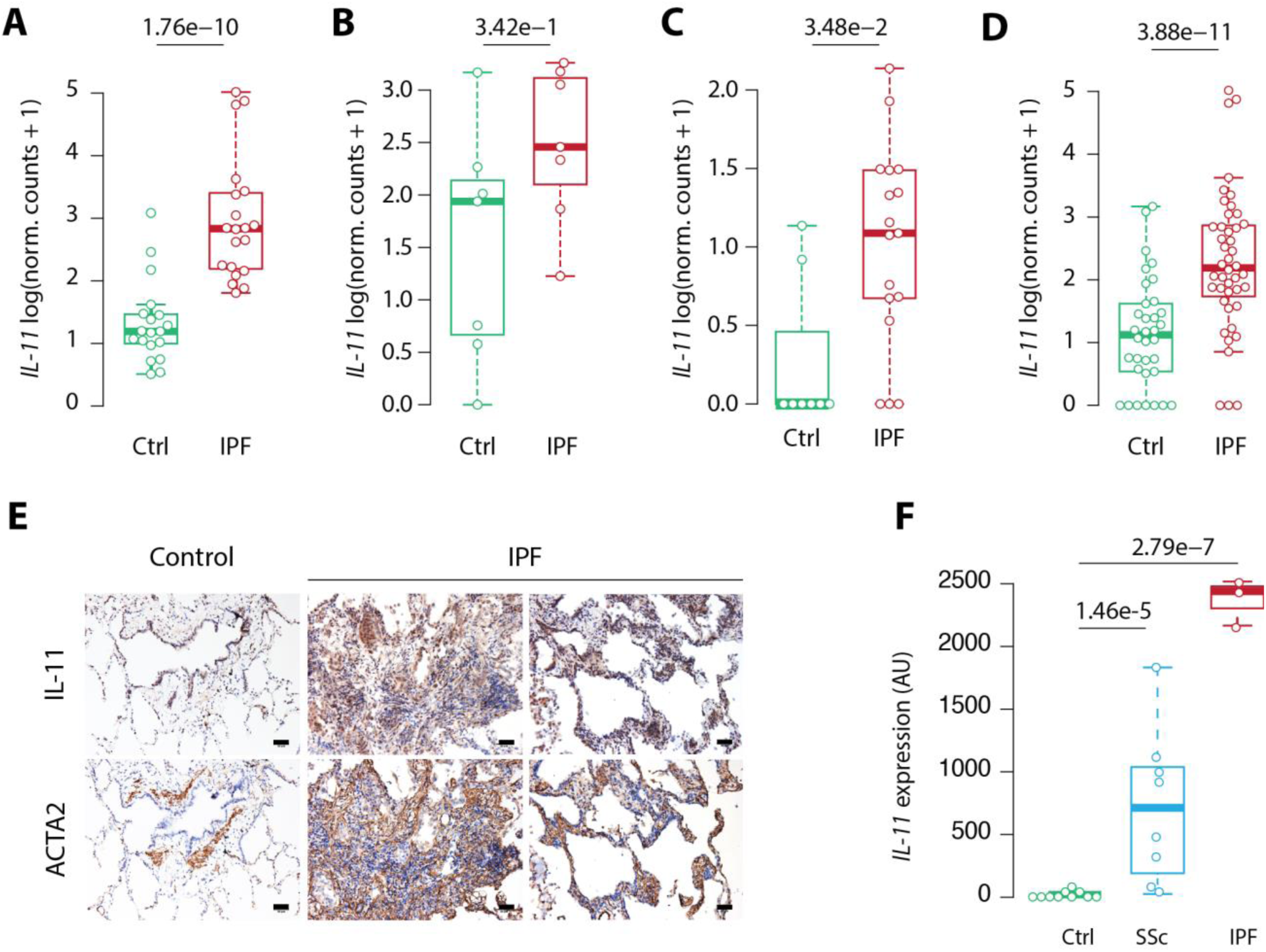
IL-11 is upregulated in IPF lung and fibroblasts. (**A-D**) Re-analysis of publicly available datasets showing elevated IL-11 RNA levels in lung tissue from IPF individuals as compared to healthy controls. (**A**) Log-transformed normalized RNA-seq counts from 19 healthy donors and 20 IPF donors sequenced in Schafer et al. 2017. (**B**) Log-transformed normalized RNA-seq counts from 7 healthy donors and 7 IPF donors sequenced in Nance et al. 2014. (**C**) Log-transformed normalized RNA-seq counts from 8 healthy donors and 17 IPF donors sequenced in Luzina et al. 2016. (**D**) Log-transformed normalized RNA-seq counts from all studies pooled together as a meta-analysis. In each panel **A**-**D**, the thick line represents the median, solid lines represent the interquartile range, and dashed lines (whiskers) represent the interquartile range x 1.5. Adjusted *P* values are reported above the black line. (**E**) Representative immunostaining of IL-11 and ACTA2 in serial sections of IPF and healthy donor (Control) lung tissue, showing increased IL-11 staining in fibrotic regions of IPF lungs. Scale bars, 50 μm. (**F**) Microarray analysis of IL-11 expression in IPF fibroblasts derived from 10 healthy donors, 8 donors with systemic sclerosis and 3 donors with IPF. Adjusted *P* values for each comparison are shown above black lines. IPF: idiopathic pulmonary fibrosis. SSc: systemic sclerosis; AU: arbitrary units.

### IL-11 is a pro-fibrotic cytokine in the lung

The IL-11 receptor subunit alpha (IL11RA) is most highly expressed in fibroblasts (*11*). To investigate the effect of IL-11 on fibrosis in the lung, we first incubated murine pulmonary fibroblasts with recombinant mouse Il-11 (Il-11, 5 ng/ml, 24h). Il-11 induced marked fibroblast activation, proliferation and ECM production as measured by high content imaging (Fig. 2A). Secreted collagen was also significantly increased in Il-11 stimulated fibroblast cultures (Fig. 2B). Fibroblast migration and invasion, critical in the pathobiology of IPF, were also driven by Il-11 (Fig. 2, C and D). Recombinant human IL-11 had identical effects on primary human pulmonary lung fibroblasts, confirming the pro-fibrotic activity of IL-11 on pulmonary fibroblasts across species (fig. S2, A to E).

**Fig. 2.**
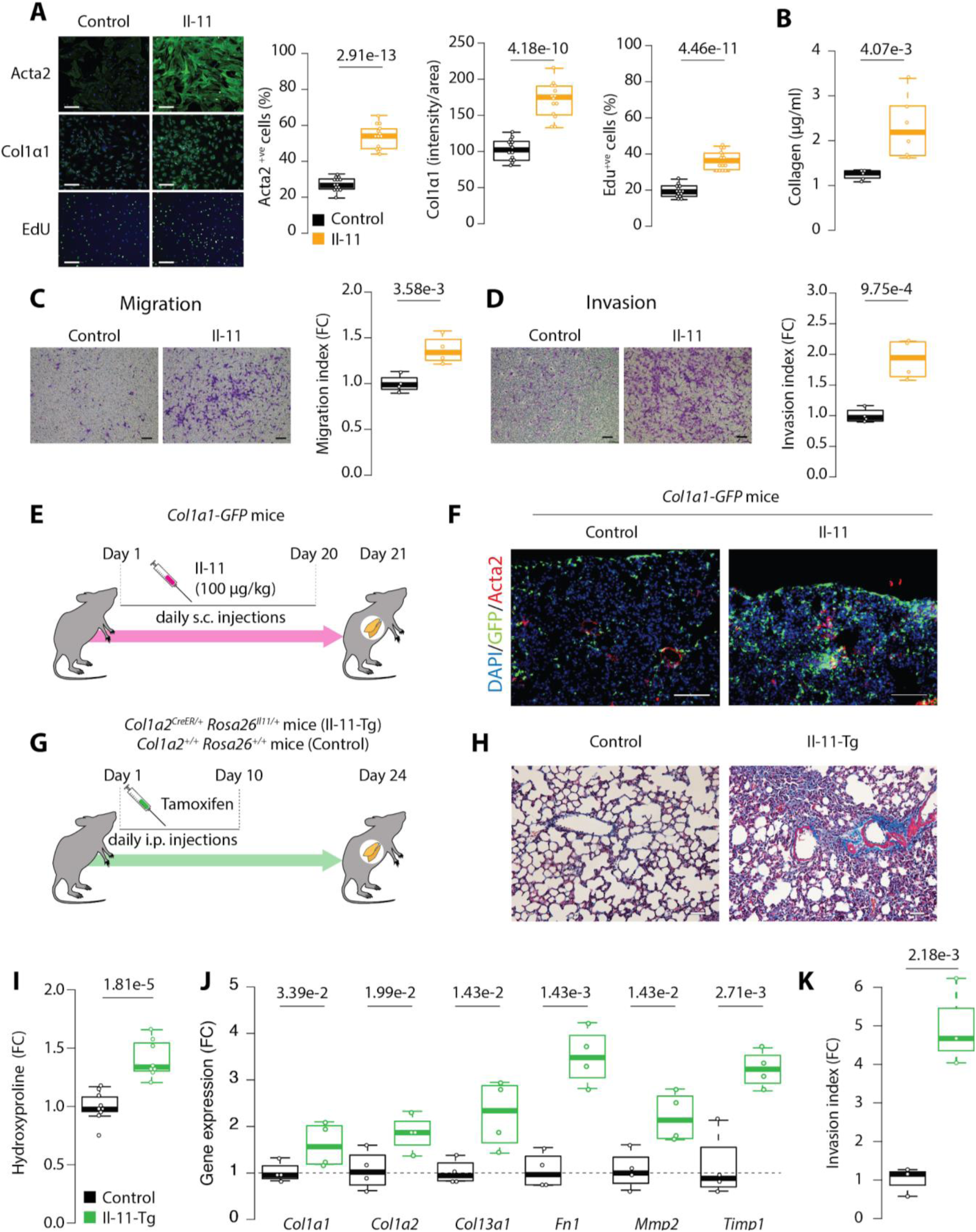
IL-11 activates lung fibroblasts *in vitro and in vivo*. (**A**) Representative immunofluorescence images (left) and quantification (right) of primary mouse lung fibroblasts stained for Acta2, Col1α1 or EdU either with or without Il-11 treatment (5 ng/ml, 24h). Cells were counterstained with DAPI to visualize nuclei. Data shown are a representative example of three independent experiments. Scale bars, 200 μm. (**B**) Total secreted collagen in the supernatant of Il-11 treated fibroblasts was quantified by Sirius red collagen assay. (**C**) Transwell migration and (**D**) matrigel invasion of wild-type mouse lung fibroblasts towards media containing Il-11 were determined. Scale bars, 150 μm. (**E**) Schematic and (**F**) representative fluorescence images of *GFP*-positive cells in lung tissue of *Col1a1-GFP* mice treated with daily subcutaneous injections of Il-11 (100 μg/kg) as compared to saline-injected controls. Sections were immunostained for Acta2 to visualize pulmonary smooth muscle cells and myofibroblasts and counterstained with DAPI to visualize nuclei. Scale bars, 100 μm. (**G**) Schematic strategy for the inducible expression of Il-11 in *Col1a2*-expressing cells in mice (Il-11-Tg). (**H**) Representative Masson’s trichrome staining of lung sections from Il-11-Tg and littermate control mice. Scale bars, 50 μm. (**I**) Relative lung hydroxyproline content and (**J**) mRNA expression of fibrosis genes in Il-11-Tg and littermate control mice lungs. (**K**) The spontaneous matrigel-invading capacity of lung fibroblasts isolated from Il-11-Tg and control mice was determined. All comparisons analysed by Student’s *t*-test; *P* values in **J** are corrected for false discovery rate using the Benjamini-Hochberg method. Data represented as median ± IQR, whiskers represent IQR x 1.5. FC: fold change.

Subcutaneous injection of Il-11 (100 µg/kg, 20d) into *Col1a1-GFP* reporter mice (*19*) resulted in an accumulation of *Col1a1*^+ve^ fibroblasts throughout the lung parenchyma (Fig. 2, E and F and fig. S3A). Administration of Il-11 also increased pulmonary collagen content and induced a fibrogenic gene expression signature in the lung (fig. S3, B and C). To model secretion of endogenous Il-11 by fibroblasts *in vivo*, we induced expression of recombinant mouse Il-11 in *Col1a2-Cre* transgenic Il-11(Il-11-Tg) mice (*11*). After 3 weeks of fibroblast-specific Il-11 expression, we observed extensive lung fibrosis, collagen deposition and an upregulation of fibrosis-related genes (Fig. 2, G to J and fig. S4A). Lung fibroblasts isolated from tamoxifen treated Il-11-Tg mice were extremely invasive, a typical feature of lung fibroblasts in IPF (*20*), in the absence of additional stimulation (Fig. 2K and fig. S4B). Taken together, these results show that activation of Il-11 signalling alone is sufficient to drive pulmonary fibrosis through activation of fibroblasts both *in vitro* and *in vivo*.

### Autocrine IL-11 signaling is required for lung fibroblast activation

The TGFβ1-driven transcriptional response in fibroblasts via SMAD signalling is a central feature of IPF (*9*). We stimulated primary human lung fibroblasts with human recombinant TGFβ1 (5 ng/ml, 24h) and performed transcriptome profiling. IL-11 was highly upregulated (fold change = 31.15, Padj.= 4.13e-97), indicating that TGFβ1 induces the fibrogenic autocrine loop of IL-11 signalling in pulmonary fibroblasts (Fig. 3A). We confirmed the upregulation of Il-11 downstream of Tgfβ1 at the protein level (Fig. 3B). In addition to TGFβ1 various other factors, including Fgf2, Pdgf, Osm, Il-13 and Endothelin 1(End1), driver fibrosis in IPF (*21*–*25*) and we found that all these pro-fibrotic stimuli induced Il-11 secretion from lung fibroblasts (Fig. 3B). This shows that Il-11 is a point of signalling convergence for diverse pro-fibrotic stimuli in lung fibroblasts. Intriguingly, Pdgf that is of particular importance in IPF and targeted by nintedanib is the most potent stimulator of Il-11 secretion.

**Fig. 3.**
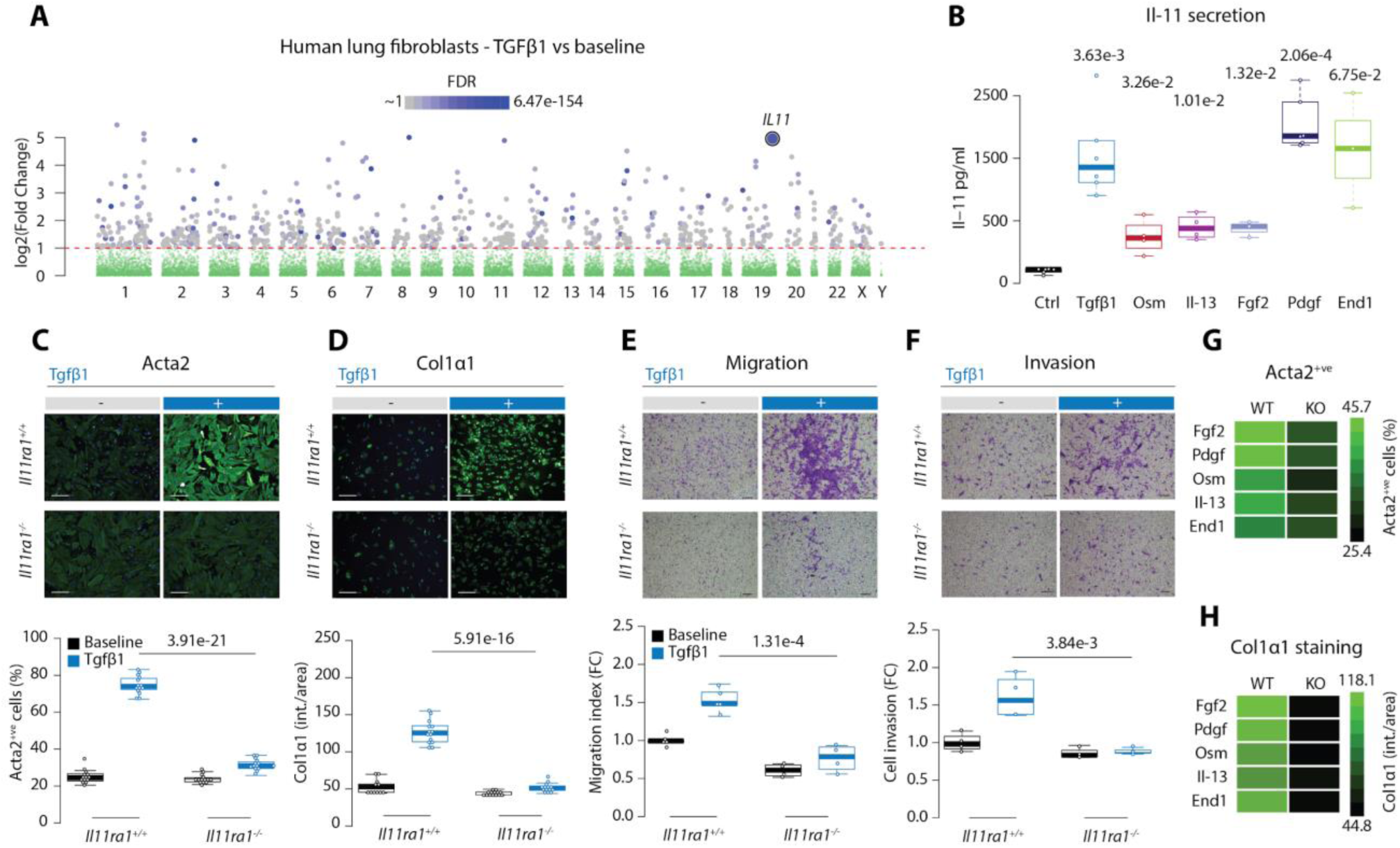
IL-11 directs lung fibroblast activation downstream of multiple pro-fibrotic stimuli. (**A**) Graphical representation of RNA-seq data showing genes upregulated in human lung fibroblasts following stimulation with TGFβ1 (5 ng/ml, 24 h). Significance values are shown only for genes with log2 (fold change) > 1 (red dashed line). FDR, false discovery rate. (**B**) ELISA of secreted Il-11 from mouse lung fibroblasts after 24h treatment with Tgfβ1 (5 ng/ml), Fgf2 (10 ng/ml), Pdgf (50 ng/ml), Osm (50 ng/ml), Il-13 (50 ng/ml) or End1 (250 ng/ml). (**C-D**) Representative immunofluorescence images (top panels) and quantification (bottom panels) of Acta2^+ve^ cells and Col1α1 immunostaining of primary lung fibroblasts from *Il11ra1*^*+/+*^ and *Il11ra1*^*-/-*^ mice treated with Tgfβ1 (5 ng/ml, 24h). Data shown are a representative example of three independent experiments. Scale bars, 200 μm. (**E-F**) Comparison of Tgfβ1-induced migration or matrigel invasion between primary lung fibroblasts from *Il11ra1*^*+/+*^ and *Il11ra1*^*-/-*^ mice. Top panels, representative images; bottom panels, quantification. Scale bars, 150 μm. (**G-H**) Quantification of Acta2^+ve^ cells and Col1α1 immunostaining (intensity/area) in *Il11ra1*^*+/+*^ and *Il11ra1*^*-/-*^ lung fibroblasts treated after Tgfβ1 stimulation (5 ng/ml, 24h). Data in **C-F** are represented as median ± IQR, whiskers represent IQR x 1.5. All comparisons analysed by Student’s *t*-test; *P* values in **B** are referred to the comparison with control values and were corrected for false discovery using the Benjamini-Hochberg procedure.

We then tested whether the various upstream pro-fibrotic stimuli were able to trigger the fibrotic response in pulmonary fibroblasts autonomously or were dependent on autocrine Il-11 signalling. Fibroblasts isolated from the lung of *Il11ra1*^*-/-*^ mice did not differentiate into myofibroblasts after Tgfβ1 stimulation as compared to those from littermate controls (Fig. 3C). Tgfβ1-induced ECM production, migration and matrigel invasion were also reduced in pulmonary fibroblasts lacking *Il11ra* (Fig. 3, D to F and fig. S5A). In addition, siRNA-mediated knockdown of *Il-11* or *Il11ra1* also reduced Tgfβ1-mediated fibroblast activation (fig. S5B). Furthermore, deletion of *Il11ra1* blocked fibroblast activation and collagen production induced by Fgf2, Pdgf, Osm, Il-13 or End 1 (Fig. 3, G and H and fig. S5, C to F). These data, across a range of approaches, show that pulmonary fibroblasts are critically dependent on autocrine IL-11 signaling if they are to become activated myofibroblasts.

### Development of a therapeutic IL-11 neutralising antibody

To translate our findings and block the pro-fibrotic IL-11 autocrine loop *in vivo*, we initiated a program to develop neutralising anti-IL-11 antibodies. Five mice were genetically immunized and 2 animals developed a strong immune response to human IL-11. Initial screenings of early hybridoma supernatants identified 16 pools of which 12 showed specific binding (Fig. 4A and fig. S6A). After further subcloning, specific binders were used in *in vitro* primary cell-based fibrosis screen to identify neutralising antibodies (Methods). Primary fibroblasts were incubated with TGFβ1 (5 ng/ml, 24h) to induce endogenous IL-11 secretion and clones were then screened for neutralisation activity using fibrosis readouts based on high-content imaging. The clone X203 most effectively blocked the fibrotic response across species and was selected for preclinical testing in mouse models of IPF and for humanisation for clinical use (Fig. 4B). Injected [^125^I]X203 had a half-life of 222h in the blood of wild type C57BL/6 mice (Fig. 4C) and the equilibrium dissociation constant (KD) for X203 was found to be in the low nM range (Fig. 4D and fig. S6B).

**Fig. 4.**
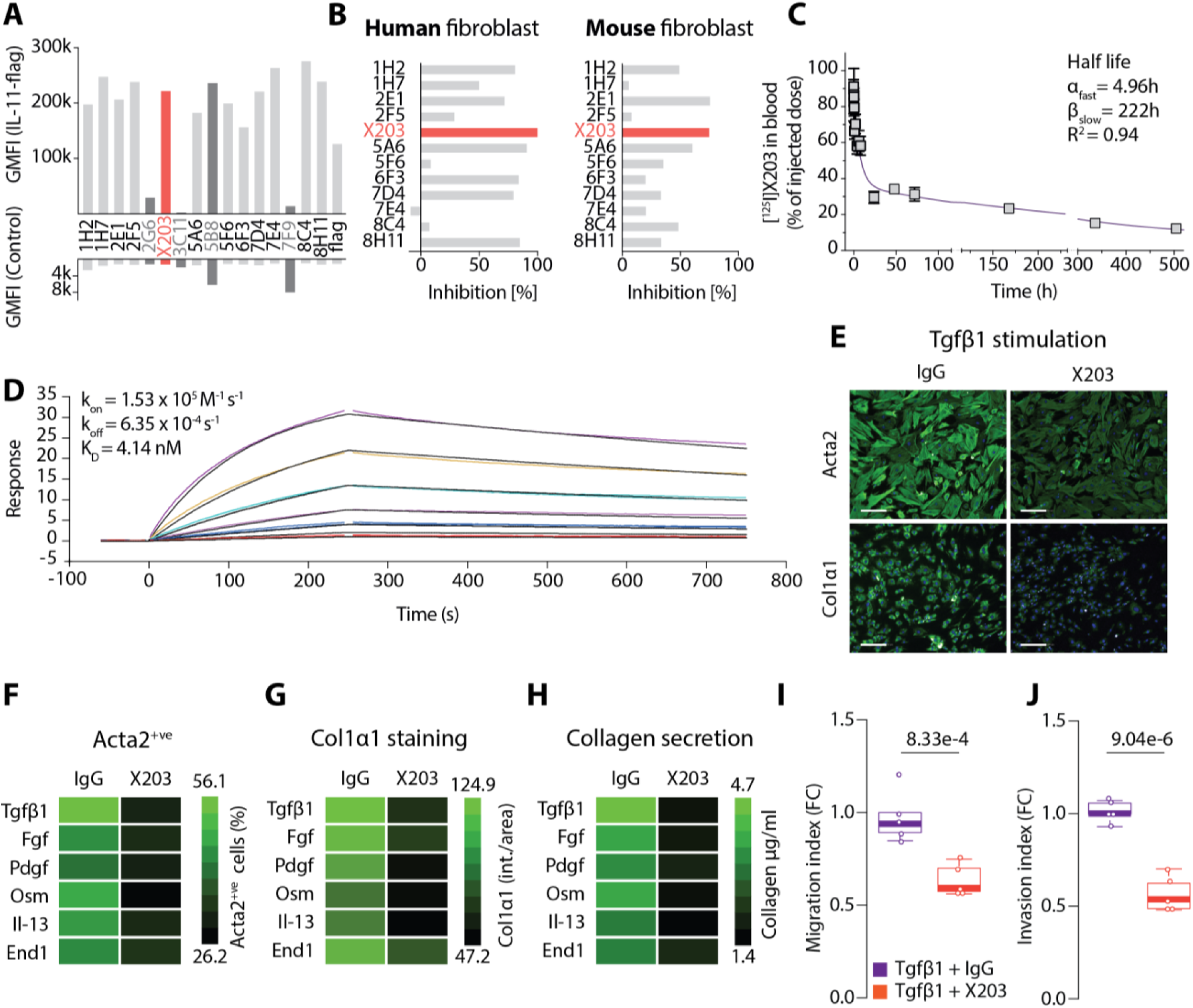
Development of therapeutic IL-11 neutralizing antibodies. (**A**) Supernatants of early stage hybridoma cultures on HEK293 cells transiently expressing IL-11-flag or an unrelated control cDNA. Signal is geometric mean of the relative fluorescence (GMFI) as measured by flow cytometry with goat anti-mouse fluorescent antibody (10 µg/ml). Non-specific binding (dark grey) and X203 (red) clones are highlighted. Positive control: anti-flag mouse antibodies. (**B**) Human and mouse primary fibroblasts were stimulated with corresponding recombinant TGFβ1 (5 ng/ml, 24h) and ACTA2^+ve^ cells were monitored in the presence of purified mouse monoclonal antibodies (6 µg/ml). X203 effectively inhibits TGFβ1-driven fibroblast activation by neutralizing the downstream IL-11 autocrine loop. (**C**) Blood pharmacokinetics of [^125^I]X203 in C57BL/6 male mice (n=5). Data are mean ± SD. Result can be fitted (R^2^=0.92) to a two-phase exponential decay model with a distribution half-life of α _fast_=4.96h and a elimination half-life of β_slow_=222h (Mouse 3, 10 min was removed from analysis). (**D**) Real-time binding kinetics of human IL-11 to X203 immobilized on a biosensor show a dissociation constant K_D_ of 4.14 nM using the BIAcore T200 system. (**E**) Representative images of Acta2 and Col1α1 immunostaining in mouse lung fibroblasts treated with Tgfβ1 (5 ng/ml, 24h) in the presence of X203 or IgG control antibodies (2 μg/ml). Scale bars, 200 μm. (**F**-**G**) Heatmaps showing the immunofluorescence quantification of Acta2^+ve^ cells and Col1α1 immunostaining (intensity/area) in fibroblasts treated with multiple pro-fibrotic stimuli in the presence of X203 or IgG control antibodies. (**H**) Collagen secretion in culture supernatant from lung fibroblasts treated as depicted in **F** and **G**. (**I**-**J**) The effects of X203 on Tgfβ1-induced migration or matrigel invasion of mouse lung fibroblasts. Data are represented as median ± IQR, whiskers represent IQR x 1.5. *P* value was determined by Student’s *t*-test.

The clone X203 effectively blocks Tgfβ1-driven fibroblast activation and ECM production (Fig 4E). Given that the Il-11 autocrine loop is also required for multiple pro-fibrotic stimuli (Fig. 3), we tested whether the X203 antibody could also block the fibrotic response downstream of Fgf, Pdgf, Osm, Il-13 or End1 stimulation. In response to all stimuli tested, pulmonary fibroblast activation, ECM secretion, migration and invasion was effectively abrogated using the X203 monoclonal antibody (Fig. 4, F to J and fig. S7, A to E). Incubation with X203 also significantly inhibited multiple fibrotic phenotypes of IPF patient-derived lung fibroblasts after TGFβ1 stimulation (fig. S7, F to H). These data show that X203 specifically binds and neutralises IL-11 and stops the fibrotic response at a central node of signalling *in vitro*.

### Therapeutic targeting of Il-11 prevents fibrosis in a mouse model of IPF

We used the established mouse model of bleomycin (BLM)-induced lung fibrosis and observed that Il-11 was largely upregulated (Fig. 5A), as documented in human IPF lung (Fig. 1). Western blot analysis showed progressive Il-11 upregulation after BLM treatment that was mirrored by increased collagen expression and activation of ERK signaling (Fig. 5B and fig. S8A). Approximately 10% of the injected dose of anti-IL-11 antibody X203 was taken up by the lung within one hour and then dropped to ∼5% after one day (Fig. 5C).

**Fig. 5.**
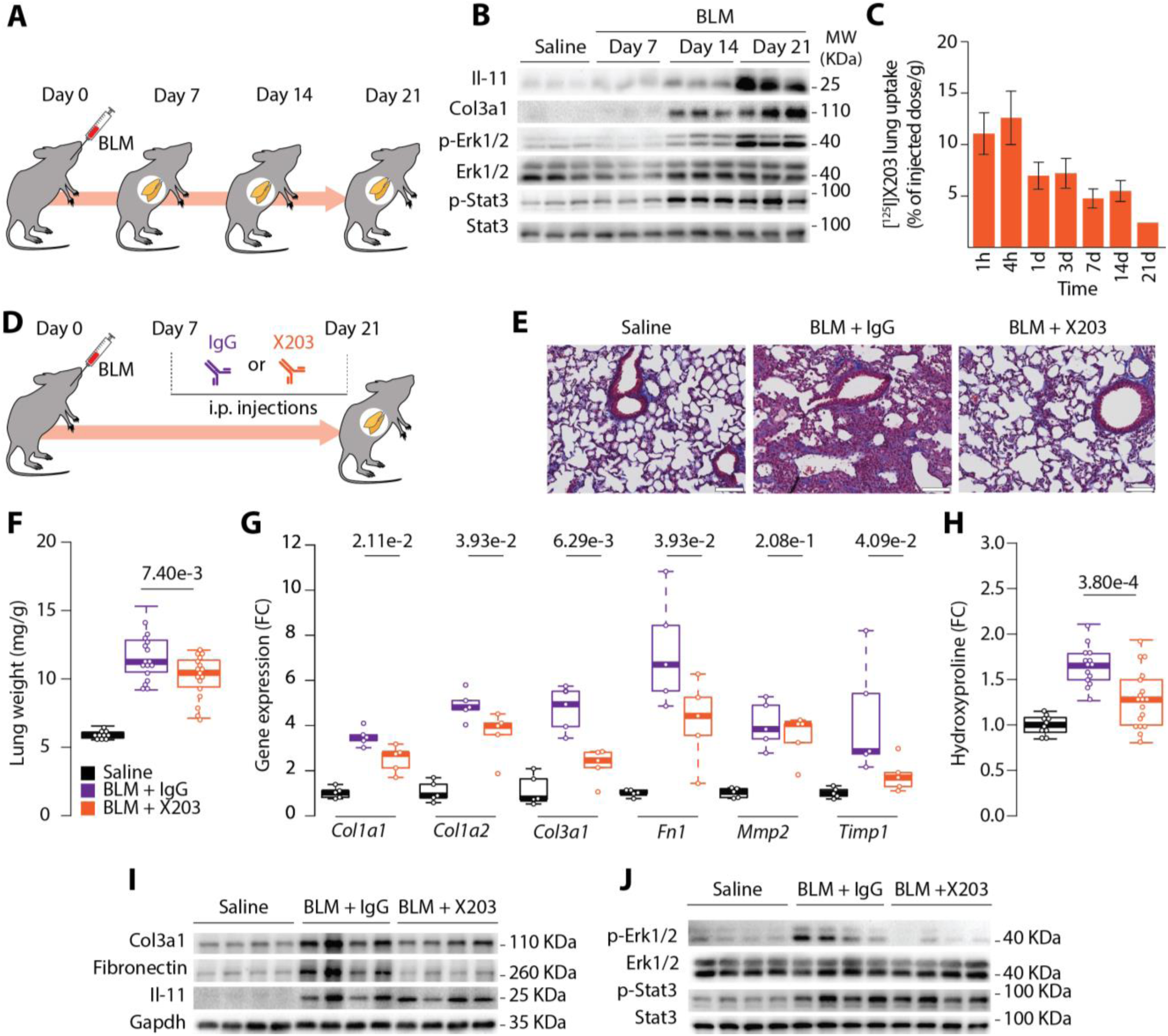
Neutralizing anti-IL-11 antibodies prevent lung fibrosis in mice. (**A**) Schematic showing the induction of lung fibrosis in mice using bleomycin (BLM). A single dose of bleomycin was injected intratracheally and lung tissue were collected at the indicated time points. (**B**) Western blot of Il-11 protein, Col3a1 and phosphorylated expression of Erk and Stat3 in total lung homogenates of bleomycin-challenged mice. (**C**) Lung tissue uptake of [^125^I]X203 in C57BL/6 mice (n=5/group). Tissue harvested at indicated time points after i.v. injection (2.5 µg/mouse). Data are mean ± SEM. (**D**) Schematic showing therapeutic blockade of Il-11 in the bleomycin lung fibrosis model. X203 (20 mg/kg, alternate days) was injected intraperitoneally starting at Day 7 post-BLM administration. (**E**) Representative Masson’s trichrome staining of lungs sections from X203- or IgG-treated mice harvested on day 21 after bleomycin-challenge. Scale bars, 50μm. (**F**) Indexed lung weights, (**G**) mRNA expression of fibrosis genes and (**H**) hydroxyproline content in the lungs of X203 or IgG antibody treated mice. (**I**) Western blots of Col3a1, fibronectin, Il-11 and Gapdh protein levels and (**J**) phosphorylation status and total levels of Erk and Stat3 in lung homogenates of X203 or IgG antibody treated mice. Data in **F, G** and **H** are represented as median ± IQR, whiskers represent IQR x 1.5. *P* values were determined by student’s *t*-test. In **H**, *P*-values are corrected for false discovery using the Benjamini-Hochberg procedure.

To investigate potential therapeutic benefits of blocking Il-11 signaling in the lung, we injected mice with either the X203 or an isotype control IgG (20 mg/kg, alternate days) from day 7 after BLM treatment (Fig. 5D). X203 treatment limited lung scarring, infiltration and parenchymal disruption and reduced lung weights as compared to the control group (Fig. 5, E and F and fig. S9). Pulmonary expression of fibrosis marker RNAs and proteins, including collagen levels using two different assays, were significantly reduced in X203-injected mice when compared to IgG control injected animals (Fig. 5, G to I and fig. S10A). Both Stat3 and Erk were activated in the lung following BLM treatment, Erk activation most closely following the time course of Il-11 upregulation (Fig. 5B and fig. S8B). Blocking Il-11 in the lung prevented Erk activation whereas Stat3 phosphorylation remained high (Fig. 5J and fig. S10B). This shows that Stat3 activation in the BLM model is independent of Il-11 activity and that the contribution of Stat3 to lung fibrosis is limited. In contrast, the activation of Erk signalling in the lung is dependent on Il-11 signaling and the action of anti-IL-11 therapies on ERK inhibition may be a useful target engagement assay.

## Discussion

There is limited and contradictory literature on the role of IL-11 in the lung (*12*–*14*, *26*, *27*) that has hindered our understanding of the function of this rarely studied cytokine in pulmonary biology. During our studies of IL-11 in the heart (*11*) we came across the provocative, chance finding that *IL-11* was the most highly upregulated gene in lung fibroblasts from IPF patients (*15*). This prompted us to study IL-11 in lung fibrosis and we show here that IL-11 is highly upregulated in the IPF lung where IL-11 drives pulmonary fibroblast activation, invasion, ECM deposition and tissue disruption via non-canonical ERK signalling. Autocrine IL-11 activity in lung fibroblasts is essential for the pro-fibrotic activity of various stimuli. This suggests that therapeutic inhibition of IL-11 at this point of signaling convergence will be more effective than blocking individual factors (e.g. IL-13 or CTGF (*8*)) for preventing lung fibrosis.

To translate our findings towards clinical application, we developed a monoclonal antibody (X203) that binds to both human and mouse Il-11 with single-digit nM affinities. By neutralising IL-11 signalling, X203 effectively blocks the fibrotic response of lung fibroblasts across species and downstream of many pro-fibrotic stimuli important in IPF. In a preclinical model of IPF, administration of X203 during the course of disease reduces ECM production and tissue disruption and specifically blocks pathogenic Erk signalling in the diseased lung.

Pirfenidone and Nintedanib are approved to treat IPF but it remains a progressive and fatal disease despite drug therapy, which is itself associated with a number of side effects including diarrhoea, photosensitivity and liver dysfunction. Inhibition of IL-11 is not expected to be associated with these, or other, side effects as it is not expressed in healthy tissues (*11*) and observations of knockout mice and humans suggest IL-11 function in healthy adult mammals is largely redundant (*28*).

We propose IL-11 as a novel drug target in IPF and believe that clinical trials of anti-IL-11 therapies in IPF, as a monotherapy or in combination with existing drugs, should be performed when such drugs are approved for use in humans.

## Acknowledgements

The authors would like to acknowledge the technical expertise and support of B.L.George, E.Khin, N.E.Sahib, M.Wang, S-Y.Lim, C-J.Pua and B-Y.Soh.

## Author contributions

B.N., S.S. and S.A.C. conceived and designed the study. B.N., J.D., S.V., A.A.W., J.T. and B.H. performed *in vitro* cell culture, cell biology and molecular biology experiments. B.N., J.D., A.A.W., N.S.K. and J.T. performed *in vivo* gain- and loss-of function mouse studies. N.G-C. and S.M.E. performed gain-of function studies on *Col1a1-GFP* mice. B.N., W-W.L., X.C. and A.J.B. performed histology analysis. A-M.C. analysed pharmacokinetics and biodistribution studies. S.V. and A.A.W performed *in vitro* antibody screening. G.D. performed computational analysis. A.J.B., T.M.M., J.L. and P.W.N. provided critical reagents for the study. B.N., J.D., S.V., G.D., A.A.W., S.P.C., S.S. and S.A.C. analyzed the data. B.N., G.D., S.S., and S.A.C prepared the manuscript with input from co-authors.

## Data and materials availability

High-throughput sequencing data generated for this study can be downloaded from the Gene Expression Omnibus (GEO) repository (data currently under submission). All other data are in the manuscript or the supplementary materials.

## Material and methods

### Animal models

Animal procedures were performed in accordance with the SingHealth Institutional Animal Care and Use Committee (IACUC) or in accordance with local guidelines from collaborating laboratories. All mice were maintained in a specific pathogen-free (SPF) environment and provided food and water *ad libitum*.

#### In vivo IL-11 administration model

Ten-week old transgenic *Col1a1-GFP* reporter mice (*19*) and wild-type C57BL/6 mice were subjected to daily subcutaneous injections with 100 μg/kg of recombinant mouse Il-11 or identical volume of saline for 20 consecutive days. The lungs were collected for histological and biochemical analyses 21 days after the start of treatment.

#### Il-11 transgenic (Il-11-Tg) model

Fibroblast-specific Il-11 overexpressing mice were on the C57BL/6 background and generated as previously described (*11*). Heterozygous *Rosa26-Il-11* mice were crossed with *Col1a2-CreER* mice (*29*) to generate double heterozygous *Col1a2-CreER:Rosa26-Il-11* mice (Il-11-Tg mice). For Cre-mediated Il-11 transgene induction, Il-11-Tg mice were injected with 50 mg/kg Tamoxifen (Sigma-Aldrich) intraperitoneally at 6 weeks of age for 10 consecutive days and the animals were sacrificed on day 21. Wild-type littermates were injected with an equivalent dose of tamoxifen for 10 consecutive days as controls.

#### Mouse model of pulmonary fibrosis

Eight to 10 week old female C57BL/6 mice were anesthetized by Isoflurane inhalation and bleomycin (Sigma-Aldrich) was injected intratracheally at a dose of 1 mg/kg body weight or saline as control. Mice were euthanized at the indicated time points and lungs were harvested for histological, biochemical analysis and transcriptome profiling.

### Antibodies and reagents

Antibodies: Anti-IL11 (PA5-36544, ThermoFisher Scientific), Anti-ACTA2 (ab7817 and ab5694, abcam), anti-COL1A1 (ab34710, abcam), anti-COL3A1 antibody (sc-271249, Santa Cruz), anti-Fibronectin antibody (ab413, Abcam), anti-p-ERK1/2 (4370, Cell Signaling), anti-ERK1/2 (4695, Cell Signaling), anti-p-STAT3 (4113, Cell Signaling), anti-STAT3 (4904, Cell Signaling) and anti-GAPDH (2118, Cell Signaling). Recombinant proteins: IL-11 (PHC0115, Life Technologies), TGFβ1 (7666-MB, R&D Systems), FGF2 (3139-FB, R&D Systems), OSM (495-MO, R&D Systems), IL13 (413-ML, R&D Systems), PDGF (220-BB, R&D Systems) and Endothelin 1 (1160/100U, Tocris). Recombinant mouse IL-11 (UniProtKB: P47873) was synthesized without the signal peptide (GenScript).

### Primary lung fibroblast cultures

Primary mouse lung fibroblasts were isolated from wild-type, Il-11-Tg or *Il11ra1*^*-/-*^ C57BL/6 mice (*30*). Lungs were minced, digested for 30 minutes in DMEM containing 100 U/ml penicillin, 100 μg/ml streptomycin and 0.14 Wunsch U/ml Liberase (Roche) with mild agitation and subsequently cultured in complete DMEM supplemented with 15% FBS, 100 U/ml penicillin and 100 μg/ml streptomycin at 37°C. Fibroblasts were explanted from the digested tissue and at 80% confluence, fibroblasts were enriched via negatively selection with magnetic beads against mouse CD45 (leukocytes), CD31 (endothelial) and CD326 (epithelial) using a QuadroMACS separator (Miltenyi Biotec) according to the manufacturer’s protocol. Mouse lung fibroblasts were used for downstream experiments between passages 3 to 5.

Human lung fibroblasts were isolated from tissue explants obtained from patients with IPF, and normal lung fibroblasts were obtained from discarded portions of normal transplant donor lung tissue as previously described (*20*, *31*, *32*). Human tissue specimens were obtained under the auspices of Cedars-Sinai Medical Centre Institutional Review Board approved protocols and the diagnosis of IPF was arrived at by standard accepted American Thoracic Society recommendations. Human lung fibroblasts were used for experiments between passages 3 to 8.

### Immunofluorescence and image analysis (Operetta)

High-content immunofluorescence imaging and quantification of fibroblast activation were performed as previously described (*11*). Briefly, lung fibroblasts were seeded in 96-well CellCarrier plates (6005550, PerkinElmer) at a density of 5 ×; 10^3^ cells per well and following experimental conditions, cells were fixed in 4% paraformaldehyde (28908, Life Technologies) and permeabilized with 0.1% Triton X-100 in PBS. EdU-Alexa Fluor 488 was incorporated using a Click-iT Edu Labelling kit (C10350, Life Technologies) according to manufacturer’s protocol. Cells were blocked using 0.5% BSA and 0.1% Tween-20 in PBS before incubation with primary antibody (anti-ACTA2 or anti-COL1A1) and visualized using Alexa Fluor 488-conjugated secondary antibody (ab150113, Abcam). Cells were counterstained with DAPI (D1306, Life Technologies) in blocking solution. Plates were scanned and images were collected with an Operetta high-content imaging system 1483 (PerkinElmer). Each condition was assayed from duplicate wells and a minimum of seven fields per well. The percentage of activated fibroblasts (ACTA2^+ve^ cells) and EdU^+ve^ cells was quantified by using Harmony software version 3.5.2 (PerkinElmer). The measurement of COL1α1 fluorescence intensity per area was done using Columbus software version 2.7.1 (PerkinElmer).

### siRNA transfection

Primary mouse lung fibroblasts were seeded in 96-well black CellCarrier (PerkinElmer) plates and transfected with 12.5 nM On-Targetplus siRNAs to *Il11* or *Il11ra1* (Dharmacon) in Opti-MEM and DMEM containing 10% FBS (ratio 1:9) using Lipofectamine RNAiMax (13778, Life Technologies) for 24h. The cells were subsequently cultured in DMEM containing 1% FBS for an additional 24h before Tgfβ1 stimulation.

### Migration and matrigel invasion assay

The motility and invasive behavior of lung fibroblasts isolated from wild-type, Il-11-Tg or *Il11ra1*^*-/-*^ mice were assays using 24-well Boyden chamber migration and invasion assays (Cell Biolabs Inc.). Fibroblasts were cultured in serum-free DMEM for 24h prior to cell migration or invasion assays. Equal numbers of fibroblast in serum-free DMEM were seeded in duplicates onto the apical chamber containing polycarbonate membranes for migration assays or onto ECM-coated matrigel for invasion assays. Fibroblasts were allowed to migrate towards Il-11 or Tgfβ1 as chemoattractants. For invasion assays, fibroblasts were allowed to invade towards DMEM containing 2% FBS. After 24h of incubation at 37°C, media was removed and non-migratory or non-invasive cells were removed using cotton swabs. The cells that migrated or invaded towards the bottom chamber were stained with cell staining solution (Cell Biolabs Inc.). Cells that migrated were colourimetrically quantified at 540 nm. Invasive cells from 5 non-overlapping fields of each membrane were imaged and counted under 40x magnification. For antibody inhibition experiments, fibroblasts were pretreated with X203 or IgG control antibodies for 15 min prior to addition of chemoattractants.

### RNA-seq

#### Generation of RNA-seq libraries

Total RNA was extracted from human and mouse lung fibroblasts using RNeasy columns (Qiagen) and quantified using Qubit RNA high sensitivity assay kit (Life Technologies). RNA integrity number (RIN) was assessed using the LabChip GX RNA Assay Reagent Kit (Perkin Elmer). TruSeq Stranded mRNA Library Preparation Kit (Illumina) was used to measure transcript abundance following manufacturer’s protocol. Briefly, 1 μg total RNA with RIN value >8 was purified by capturing mRNA with polyA+ before fragmentation and reversed transcribed into cDNA. cDNA was ligated with adaptors and amplified. All final libraries were quantified using KAPA library quantification kits (KAPA Biosystems) on StepOnePlus Real-Time PCR system (Applied Biosystems) according to manufacturer’s guide. The quality and average fragment size of the final libraries were also determined using LabChip GX DNA High Sensitivity Reagent Kit (Perkin Elmer). Libraries with unique indexes were pooled and sequenced on a NextSeq 500 benchtop sequencer (Illumina) using NextSeq 500 High Output v2 kit and paired-end 75-bp sequencing chemistry.

#### Analysis of RNA-seq datasets

In-house sequenced libraries were demultiplexed using bcl2fastq v2.19.0.316 with the *--no-lane-splitting* option. Adapter sequences were then trimmed using trimmomatic (*33*) v0.36 in paired end mode with the options *MAXINFO:35:0.5 MINLEN:35*. Trimmed reads were aligned to the *Homo sapiens* GRCh38 using STAR (*34*) v. 2.2.1 with the options *--outFilterType BySJout -- outFilterMultimapNmax 20 --alignSJoverhangMin 8 --alignSJDBoverhangMin 1 --outFilterMismatchNmax 999 --alignIntronMin 20 --alignIntronMax 1000000 -- alignMatesGapMax 1000000* in paired end, single pass mode. Only unique alignments were retained for counting. Counts were calculated at the gene level using the FeatureCounts module from subread (*35*) v. 1.5.1, with the options *-O -s 2 -J -T 8 -p -R -G*. The Ensembl release 92 hg38 GTF was used as annotation to prepare STAR indexes and for FeatureCounts.

Publicly available libraries from GEO series GSE92592, GSE52463 and GSE99621 were downloaded and integrated in the same pipeline without the demultiplexing step.

Differential expression analyses were performed in R 3.4.1 using the Bioconductor package DESeq2 (*36*) 1.18.1, using the Wald test for comparisons and including the variance shrinkage step setting a significance threshold of 0.05. In particular, for the combined analysis of public datasets, the design for the linear model was specified as condition (IPF/control) + study, to account for the confounding effect of different library preparations in different studies. For our human lung fibroblast RNA-seq analysis, the design for the model was specified as stimulus (TGFβ1/baseline) + source (commercial/patient), to account for the confounding effect of different sources of fibroblasts. Batch removal was simulated using the *combat* function in the Bioconductor package sva (*37*, *38*) v. 3.26.0 and visualized using principal component analysis. In all cases, the first component was correctly identifying maximum variance between control and IPF/stimulated samples only when simulating batch removal, thus validating the inclusion of the confounding variable in the model.

### Microarray analysis

Data from the GEO dataset GSE40839 was analyzed using the GEO2R platform, which implements the GEOquery (*39*) and limma (*40*, *41*) R packages.

### ELISA and colorimetric assays

The levels of IL-11 in equal volumes of cell culture supernatant was quantified using the mouse IL-11 DuoSet ELISA kit (DY418, R&D Systems). Secreted collagen in the cell culture supernatant was quantified using a Sirius red collagen detection kit (9062, Chondrex). Total hydroxyproline content in the right lungs (superior lobe) of mice were measured using a Quickzyme Total Collagen assay kit (Quickzyme Biosciences). All ELISA and colorimetric assays were performed according to the manufacturer’s protocol.

### RT-qPCR

Total RNA was extracted from either the snap-frozen tissues or cell lysate using Trizol reagent (Invitrogen) followed by RNeasy column (Qiagen) purification. The cDNA was prepared using an iScript cDNA synthesis kit (Biorad), in which each reaction contained 1 μg of total RNA, as per the manufacturer’s instructions. Quantitative RT–PCR gene expression analysis was performed in triplicates with TaqMan (Applied Biosystems) technology using a StepOnePlus (Applied Biosystem) over 40 cycles. Expression data were normalized to GAPDH mRNA expression levels and we used the 2^−ΔΔCt^ method to calculate the fold change. All TaqMan probes were obtained from Thermo Fisher Scientific (Col1a1, Mm00801666_g1; Col1a2, Mm00483888_m1; Col3a1, Mm01254476_m1; Fn1, Mm01256744_m1; Mmp2, Mm00439498_m1; Timp1, Mm01341361_m1; Gapdh, Mm99999915_g1).

### Immunoblotting

Western blot analysis was carried out on total protein extracts from mouse lung tissues. Frozen tissues were homogenized by gentle rocking in lysis buffer (RIPA buffer containing protease and phosphatase inhibitors (Roche)) followed by centrifugation to clear the lysate. Equal amounts of protein lysates were separated by SDS-PAGE, transferred to a PVDF membrane, and incubated overnight with primary antibodies. Proteins were visualized using the ECL detection system (Pierce) with the appropriate secondary antibodies: anti-rabbit HRP (7074, Cell Signaling) or anti-mouse HRP (7076, Cell Signaling).

### Histology

Human IPF tissues were obtained from lung transplant patients with IPF and normal control human lung tissues were obtained from IIAM (International Institute for the Advancement of Medicine). Human lung tissue were fixed in 10% formalin overnight and embedded in paraffin. For mouse models, freshly dissected lungs from *Col1a1-GFP* mice were cooled in liquid nitrogen and tissues were embedded with Tissue-Tek Optimal Cutting Temperature compound (VWR International) before being subjected to cryosectioning. In other mouse models, lungs were fixed in 10% formalin for 24h, dehydrated and embedded in paraffin. For Masson’s trichrome staining, lung sections were subjected to Bouin’s fixative, Beibrich Scarlet-Acid Fuchsin and differentiated in 5% Phosphomolybdic -phosphotungstic acid, counterstained in 2.5% Aniline blue and further differentiated in 1% Acetic acid. For immunohistochemistry, tissue sections were incubated with primary antibodies overnight and visualized using an ImmPRESS HRP anti-rabbit IgG polymer detection kit (Vector Laboratories) with ImmPACT DAB Peroxidase Substrate (Vector Laboratories).

### Generation of mouse monoclonal antibodies against IL-11 (X203)

A cDNA encoding amino acid 22-199 of human IL-11 was cloned into expression plasmids (Aldevron Freiburg GmbH, Freiburg, Germany). Mice were immunised by intradermal application of DNA-coated gold-particles using a hand-held device for particle-bombardment (“gene gun”). Cell surface expression on transiently transfected HEK cells was confirmed with anti-tag antibodies recognising a tag added to the N-terminus of the IL-11 protein. Sera were collected after 24 days and a series of immunisations and tested in flow cytometry on HEK293 cells transiently transfected with the aforementioned expression plasmids. The secondary antibody used was a goat anti-mouse IgG R-phycoerythrin-conjugated antibody (Southern Biotech, #1030-09) at a final concentration of 10 µg/ml. Sera were diluted in PBS containing 3% FBS. Interaction of the serum was compared to HEK293 cells transfected with an irrelevant cDNA. Specific reactivity was confirmed in 2 animals and antibody-producing cells were isolated and fused with mouse myeloma cells (Ag8) according to standard procedures. Hybridomas producing antibodies specific for IL-11 were identified by screening using the Operetta high content imaging of ACTA2 immunostaining as described above.

### Surface plasmon resonance (SPR)

SPR measurements were performed on a BIAcore T200 (GE Healthcare, USA) at 25°C. All SPR experiments were carried out using carboxymethylated dextran (CM5) sensor chips. Buffers were degassed and filter-sterilized through 0.2 μm filters prior to use. IL-11 was immobilized onto a CM5 sensor chip using standard amine coupling chemistry (*42*). For kinetic analysis, a concentration series of X203 (0.78 nM to 50 nM) was injected over the IL-11 and reference surfaces at a flow rate of 40 μl/min. All the analytes were in Hepes-buffered saline (HBS) pH 7.4 containing 0.005% P20. The association and dissociation were measured for 250 s and 500 s respectively. After each sample injection, the surface was regenerated by a 30 s injection of 3.8M of MgCl2), followed by a 5 min stabilisation period. All sensorgrams were aligned and double-referenced. Affinity and kinetic constants were determined by fitting the corrected sensorgrams with the 1:1 Langmuir model using BIAevaluation 3.0 software (GE Healthcare). The equilibrium binding constant KD was determined by the ratio of the binding rate constants kd/ka.

### Pharmacokinetics and Biodistribution studies

Retro orbital injections were performed on C57BL/6 male mice (n=5, 10-12 week old) with freshly labeled [^125^I]X203 (4.2 μCi/2.5 μg/100 μl) in PBS per mouse. Mice were anesthetized with 2% isoflurane and blood collected at several time points: 2, 5, 10, 15, 30 min, 2h, 6h, 8h and 2d post injection via submandibular bleeding. For biodistribution studies, blood was collected via cardiac puncture and lung tissues were harvested from mice euthanized at the following time points: 1h, 4h, 1d, 3d, 7d, 14d and 21d post injection (n=5). The collected samples were weighed and radioactivity content measured using a gamma counter (2480 Wizard^2^, Perkin Elmer) with decay-corrections to input tubes (100x dilution of 100 μl dose). The measured radioactivity was normalized to % injected dose/per gram tissue.

### Statistical analysis

Statistical analysis was performed using GraphPad Prism software (version 6.07). For Operetta high-content imaging analysis, fluorescence intensity for COL1α1 staining was normalized to the total number of cells detected in each field. Cells expressing ACTA2 were quantified and the percentage of activated fibroblasts (ACTA2^+ve^ cells) was determined in each field. Statistical analyses were performed using Student’s *t*-tests or by one-way analysis of variance (ANOVA) as indicated in the figure legends. *P* values are corrected for multiple hypothesis testing using either the Benjamini-Hochberg procedure (for experiments comparing multiple couples of control and treatment) or using Dunnett’s test (for experiments comparing multiple treatments to a single control).

**Fig. S1.**
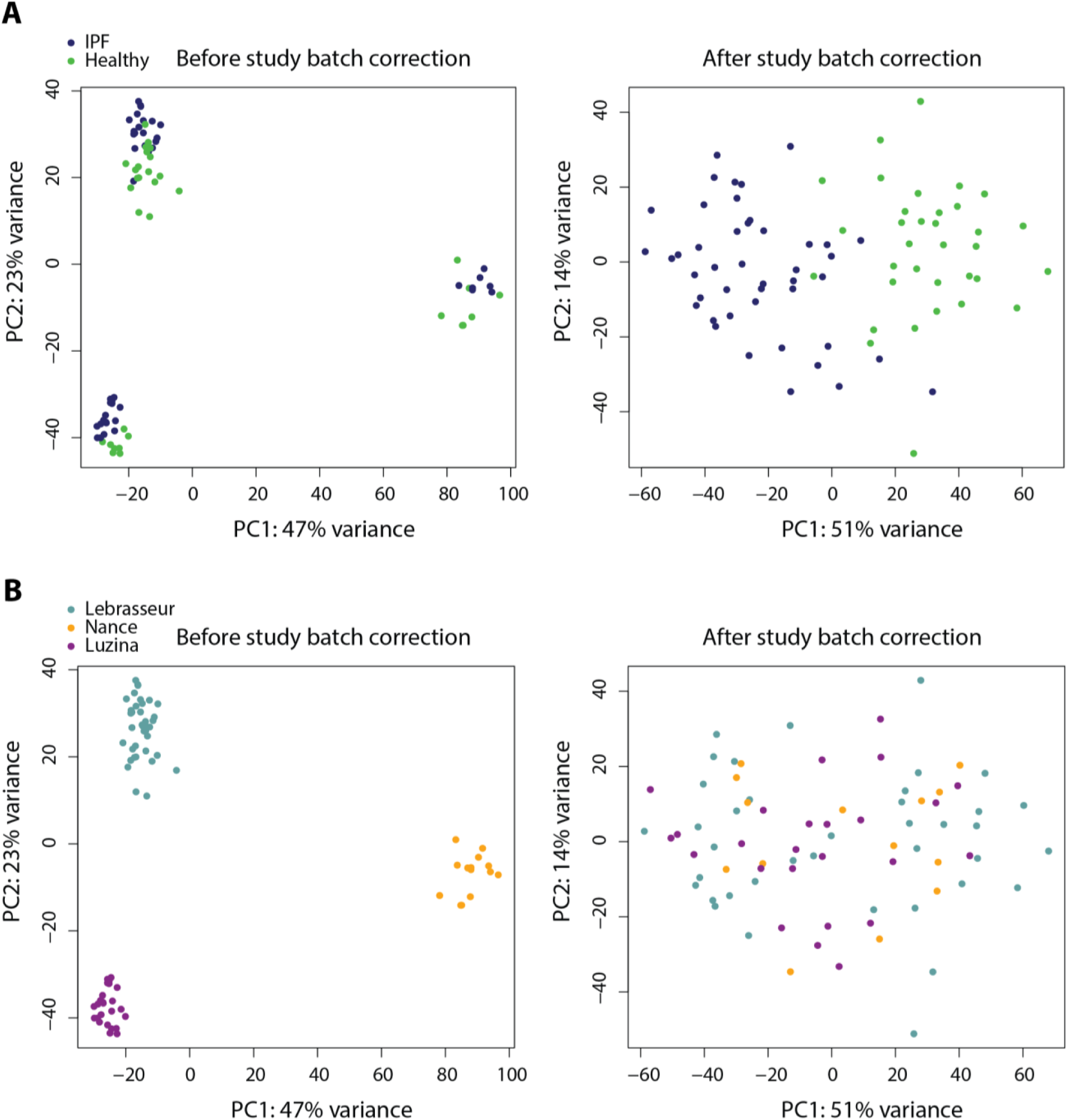
Accounting for the batch effect in the combined transcriptomic analysis identifies differences due to disease state. (**A**) Principal component analysis (PCA) of data from the three studies used in the combined analysis. PCs were determined using normalized log-transformed counts (left-hand side) and counts in which the batch effect correction has been simulated (right-hand side). Colors indicate disease state. (**B**) same as in A, but colors indicate each study. PC: principal component. IPF: idiopathic pulmonary fibrosis.

**Fig. S2.**
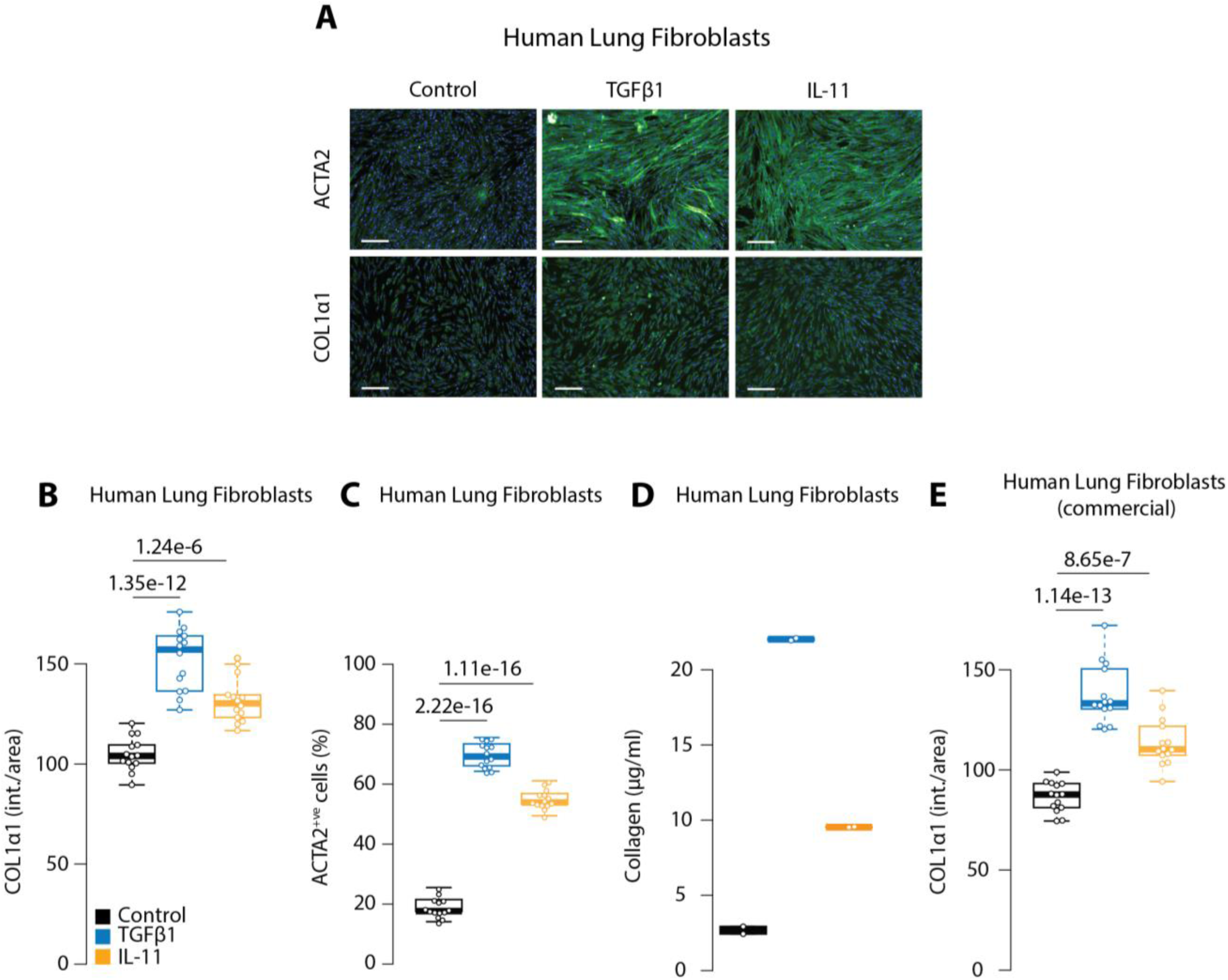
IL-11 directs human lung fibroblast activation and collagen secretion. ***in vitro.*** (**A**) Representative immunofluorescence images of primary human lung fibroblasts stained for ACTA2 or COL1α1 after 24h treatment with TGFβ1 (5 ng/ml) or IL-11 (5 ng/ml). Scale bars, 200 μm. (**B-C**) Automated quantification of fluorescence reveals significant fibroblast activation (ACTA2^+ve^) and COL1α1 expression induced by TGFβ1 or IL-11 as shown in **A**. Data shown are a representative example of three independent experiments. (**D**) Secreted collagen into the supernatant was quantified by Sirius red collagen assay (*n*=2). (**E**) Automated quantification of fluorescence of COL1α1 immunostaining in ACTA2^−ve^ human lung fibroblasts (H-6013, Cell Biologics), after treatment with TGFβ1 (5 ng/ml) or IL-11(5 ng/ml) for 24h. *P* values were determined by one-way ANOVA and corrected for comparisons to the same sample (Control) using Dunnett’s test.

**Fig. S3.**
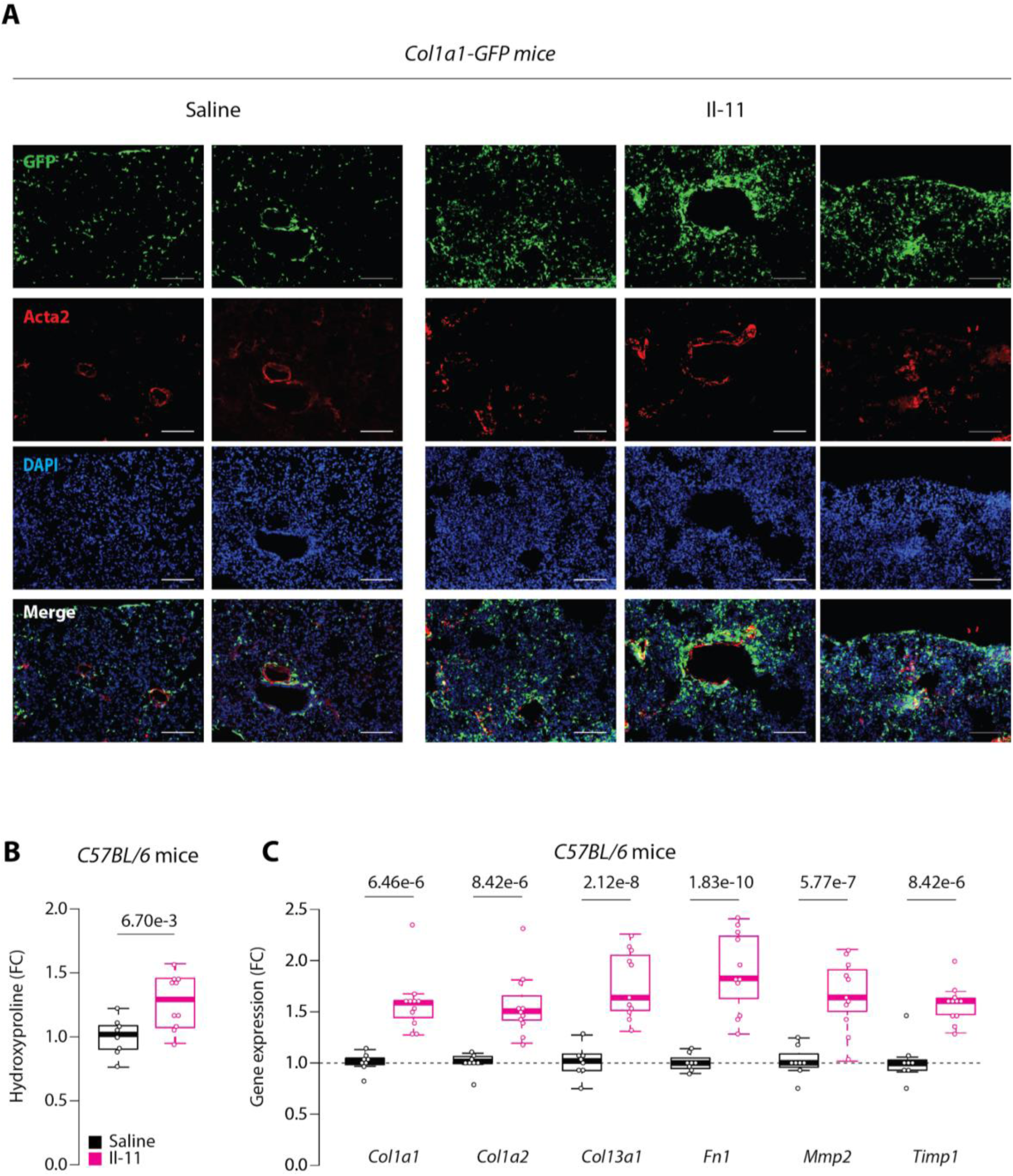
IL-11 injections in mice induces fibroblast activation and collagen deposition in the lung. (**A**) Fluorescence images of lung tissue from *Col1a1-GFP* mice treated with daily subcutaneous injections of recombinant mouse Il-11 (Il-11, 100 μg/kg, 20d) as compared to saline-injected controls. Lung sections were stained for Acta2 to visualize pulmonary smooth muscle cells, myofibroblasts and counterstained DAPI to visualize cell nuclei. Scale bars, 100 μm. (**B**) Relative hydroxyproline content and (**C**) mRNA expression of fibrosis genes in Il-11 injected as compared to saline-injected mice lungs. *P* values were determined by Student’s *t*-tests. *P* values in **C** are corrected for multiple testing using the Benjamini-Hochberg method. Data represented as median ± IQR, whiskers represent IQR x 1.5.

**Fig. S4.**
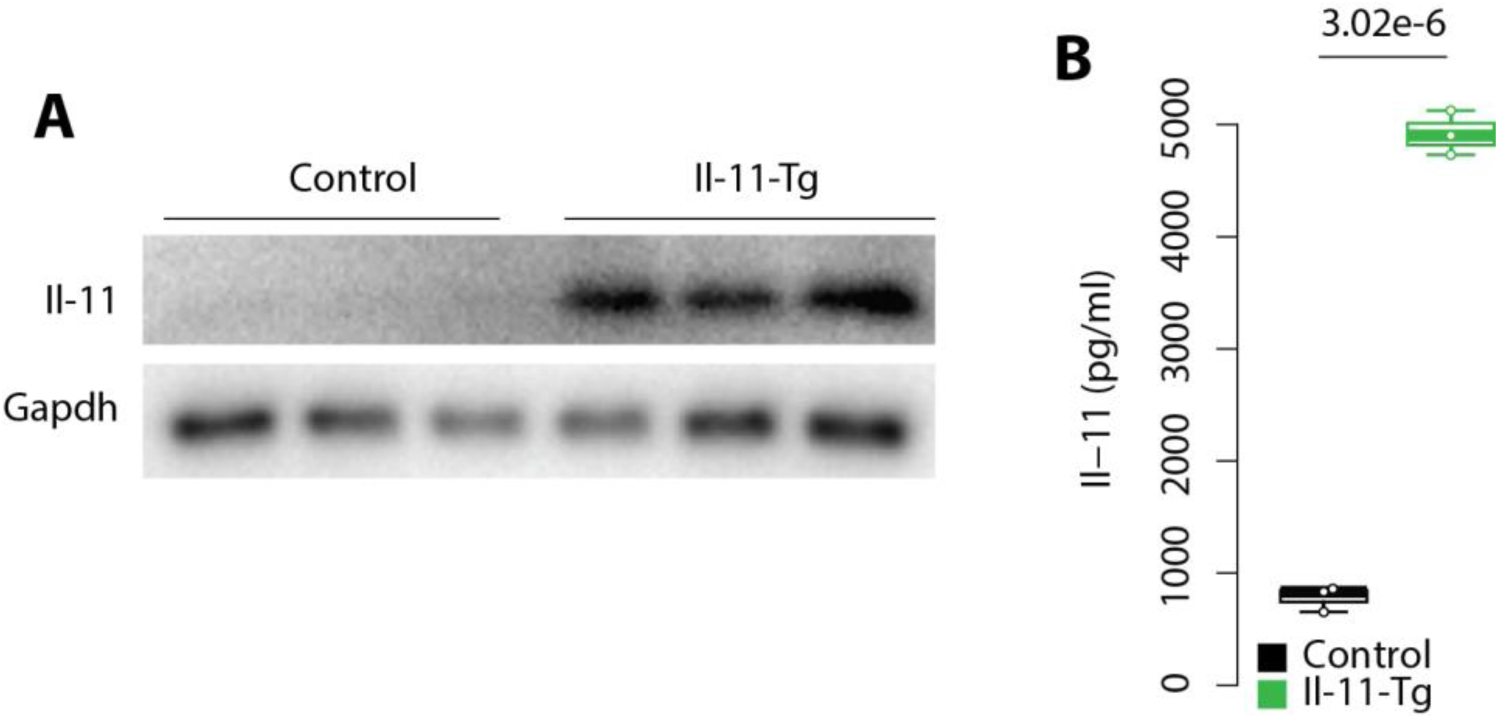
Expression of Il-11 protein in Il-11-Tg mice lung tissue and fibroblasts. (**A**) Western blot of Il-11 showing overexpression of Il-11 protein in lung tissue from tamoxifen-treated Il-11-Tg as compared to littermate control mice. (**B**) ELISA measurements of secreted Il-11 protein in the supernatant of fibroblasts isolated from Il-11-Tg and control mice.

**Fig. S5.**
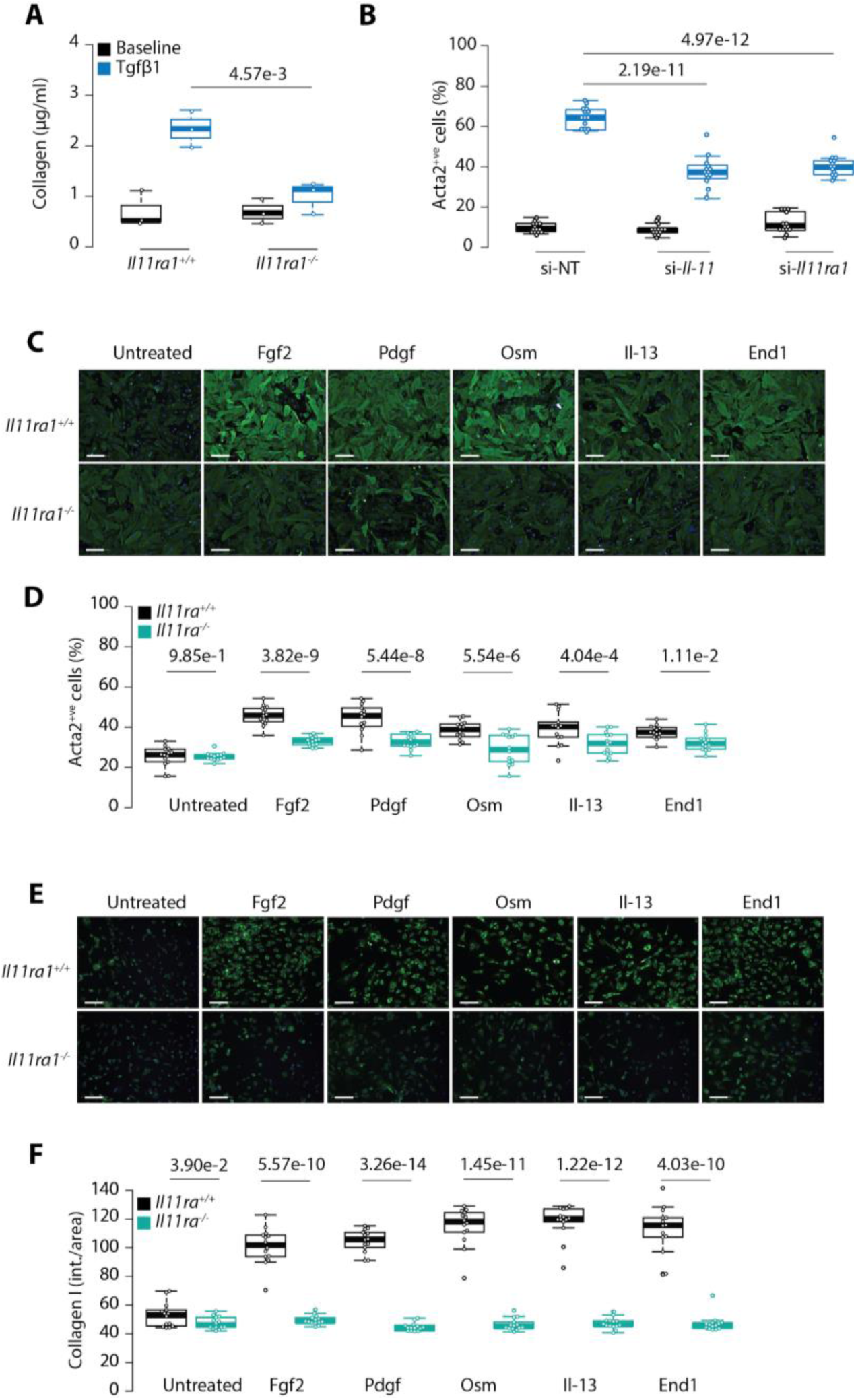
IL-11 is required for the pro-fibrotic effects of multiple stimuli in lung fibroblasts. (**A**) Secreted collagen in supernatant of *Il11ra1*^*+/+*^ and *Il11ra1*^*-/-*^ lung fibroblasts treated with Tgfβ1 (5 ng/ml, 24h). (**B**) Effects of siRNA-mediated knockdown of *Il11* or *Il11ra1* on Tgfβ1-induced myofibroblast differentiation as determined by Acta2 immunofluorescence quantification. si-NT, non-targeting siRNA. (**C**-**F**) Representative fluorescence images and quantification of (**C**-**D**) Acta2 and (**E**-**F**) Col1α1 immunostaining of lung fibroblasts from *Il11ra1*^*+/+*^ and *Il11ra1*^*-/-*^ mice treated with various pro-fibrotic stimuli for 24h, as depicted by heatmaps in Fig. 3G and 3H. Data in **D** and **F** are a representative example of three independent experiments. Scale bars, 200 μm. *P* value determined by Student’s *t-*test. *P* values in panels **B**, **D** and **F** were adjusted using the Benjamini-Hochberg method.

**Fig. S6.**
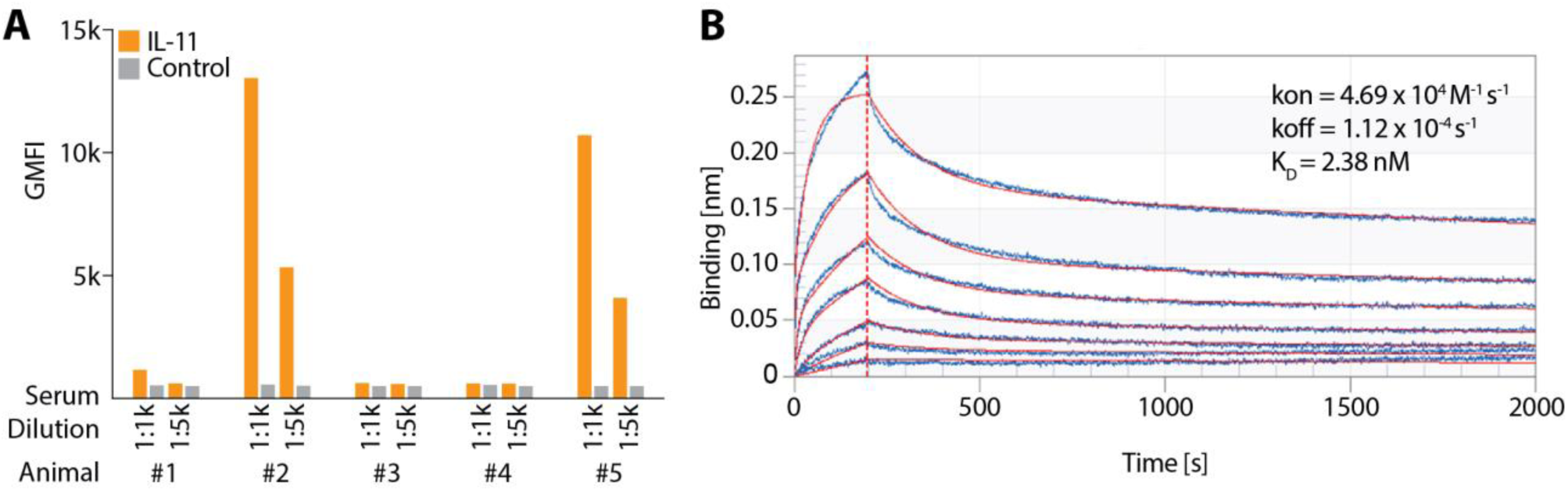
Development of neutralizing anti-IL-11 antibodies. (**A**) Sera of 5 mice after genetic immunization with human IL-11. Sera were diluted and tested with HEK cells transiently transfected with an *IL11* or control cDNA vector, incubated with a goat anti-mouse fluorescent antibody (10 µg/ml) and cells were then analysed by flow cytometry. Values represent the geometric mean of the relative fluorescence (GMFI). (**B**) Real time binding kinetics of mouse Il-11 to X203 immobilized on a Octet biosensor suggest a dissociation constant KD of 2.38 nM (2:1 model fitting).

**Fig. S7.**
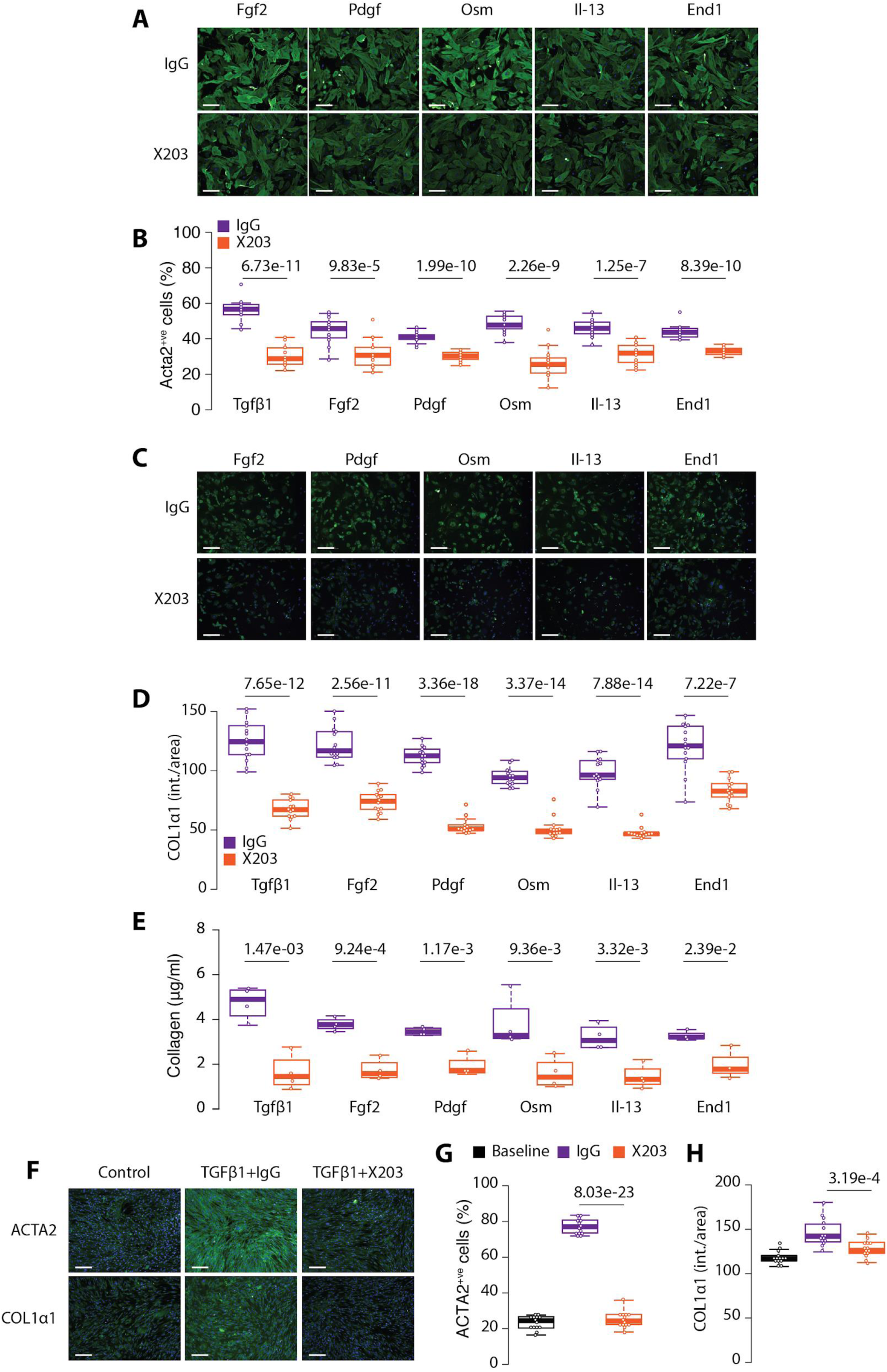
Pharmacological inhibition of IL-11 blocks fibroblast activation. Representative fluorescence images and quantification of (**A**-**B**) ACTA2^+ve^ cells and (**C**-**D**) Col1α1 immunostaining in wild-type mouse lung fibroblasts treated with various pro-fibrotic cytokines in the presence of X203 or IgG control antibodies (2 μg/ml) for 24h, as depicted by heatmaps in Fig. 4F and G. (**E**) Quantification of secreted collagen in culture media by Sirius red assay as depicted by a heatmap in Fig. 4H (*n*=3-5/group). (**F**-**H**) Representative fluorescence images and quantification of ACTA2^+ve^ cells and COL1α1 immunostaining in IPF lung fibroblasts treated with TGFβ1 (5 ng/ml, 24h) in the presence of X203 or IgG control antibodies (2 μg/ml). Data shown in **B** and **D** are a representative example of three independent experiments. Scale bars, 200 μm. *P* values were determined using Student’s *t*-test and corrected using the Benjamini-Hochberg method in panels **B**, **D** and **E**.

**Fig. S8.**
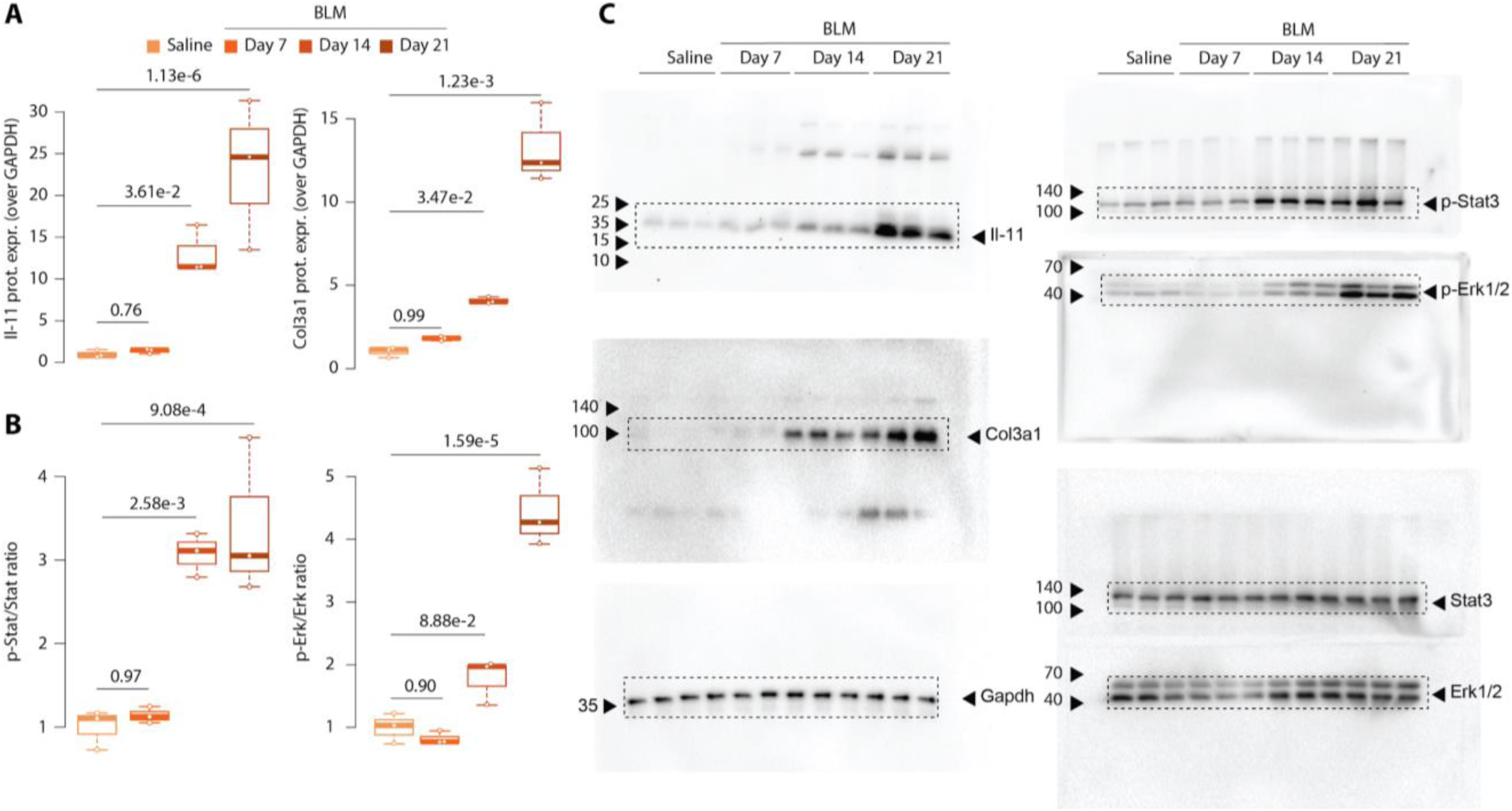
Densitometry analysis of Western blots for Fig. 5B. (**A**) Densitometry analysis for Western blots showing significant increase in Il-11 and Col3a1 protein levels in lung homogenates of bleomycin (BLM)-challenged mice. (**B**) Densitometry analysis of Western blots showing significant increase in phosphorylated Erk1/2 and Stat3 after bleomycin challenge. (**C**) Uncropped blots for Fig. 5B.

**Fig. S9.**
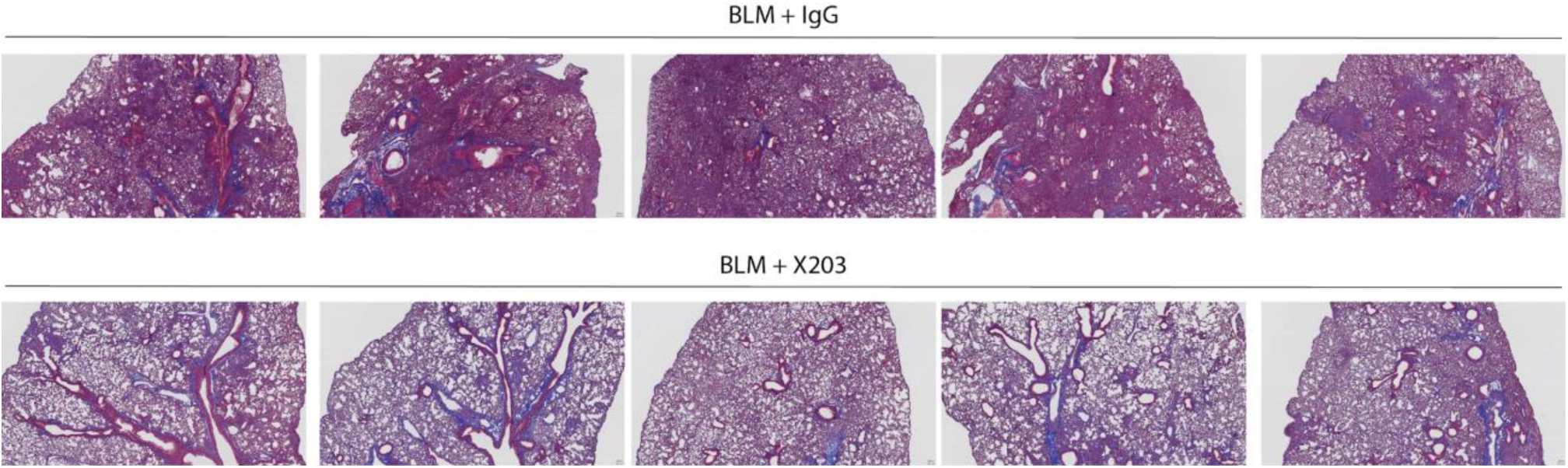
X203 treatment prevents bleomycin-induced lung damage. Masson’s trichrome staining of lungs sections from X203- or IgG-treated mice harvested on day 21 after bleomycin-challenge. Representative images of 5 mice per treatment group are shown. Images were taken at 40x magnification. Scale bars, 100 μm.

**Fig. S10.**
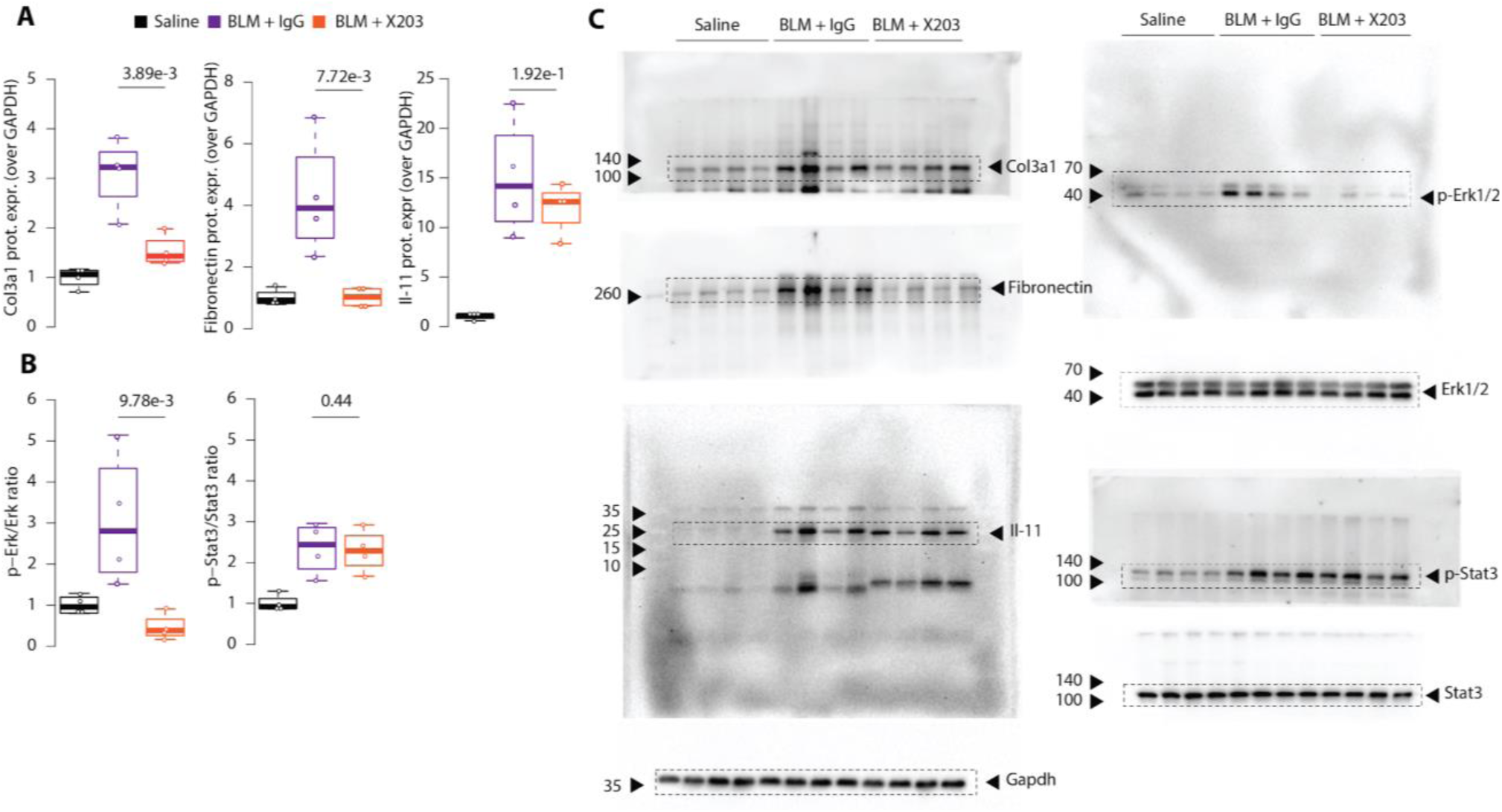
Densitometry analysis of Western blots for Fig. 5I and J. (**A**) Densitometry analysis of Western blots showing significant decrease in Col3a1 and fibronectin protein levels in lung homogenates of bleomycin (BLM)-challenged mice treated with X203- as compared to IgG-treated mice. (**B**) Densitometry analysis of Western blots showing significant decrease in phosphorylated Erk1/2, but similar levels of Stat3 phosphorylation in X203 as compared to IgG-treated mice lungs. (**C**) Uncropped blots for Fig. 5I and J.

